# AP profiling resolves co-translational folding pathway and chaperone interactions *in vivo*

**DOI:** 10.1101/2023.09.01.555749

**Authors:** Xiuqi Chen, Christian M. Kaiser

## Abstract

Natural proteins have evolved to fold robustly along specific pathways. Folding begins during synthesis, guided by interactions of the nascent protein with the ribosome and molecular chaperones. However, the timing and progression of co-translational folding remain largely elusive, in part because the process is difficult to measure in the natural environment of the cytosol. We developed a high-throughput method to quantify co-translational folding in live cells that we term Arrest Peptide profiling (AP profiling). We employed AP profiling to delineate co-translational folding for a set of GTPase domains with very similar structures, defining how topology shapes folding pathways. Genetic ablation of major nascent chain-binding chaperones resulted in localized folding changes that suggest how functional redundancies among chaperones are achieved by distinct interactions with the nascent protein. Collectively, our studies provide a window into cellular folding pathways of complex proteins and pave the way for systematic studies on nascent protein folding at unprecedented resolution and throughput.

## Introduction

Proteins are essential for all cellular functions. Commensurate with their enormous functional diversity, natural proteins exhibit a broad range of structures. Most proteins, ranging from small single-domain to large multi-domain proteins, must fold into specific three-dimensional structures to become active. In contrast to the small proteins that have principally been used as models for mechanistic folding studies^1^, multi-domain proteins exhibit a high propensity to misfold *in vitro*^2^. In the cell, folding begins during the highly regulated process of translation, while the growing nascent polypeptide emerges from the ribosome exit tunnel^3–5^. The nascent chain can adopt limited structure while still confined within the ribosome. Stable tertiary structure is mostly formed upon exiting from the tunnel. Co-translational structure formation is particularly important for efficient folding of large multi-domain proteins that are ubiquitous in all proteomes^1,6,7^.

During polypeptide elongation, all proteins interact with the ribosome, which has been shown to profoundly influence co-translational folding events^8^. Interactions with ribosomal protein and RNA residues lining the exit tunnel can stabilize structures of the elongating nascent protein^9–11^, some of which can, in turn, regulate ribosome activity by causing elongation arrest^12–14^. Once the nascent chain is extruded, ribosome surface interactions can modulate folding rates^8,15^, and influence structure stability^16^. Furthermore, molecular chaperones^17–19^ and macromolecular crowding^20^ exert effects on their stabilities. As a result, nascent chains transition through a complex set of environments. How these interactions affect co-translational folding is just beginning to be understood. Experimental studies are needed to define principles of co-translational folding.

Several experimental techniques have been employed to observe folding near the ribosome. Nuclear magnetic resonance (NMR) spectroscopy^21^ and cryogenic electron microscopy^22^ have provided detailed information on the structures of partially and fully folded domains in ribosome-nascent chain complexes. Pulse proteolysis^23^ combined with fluorescence measurements^24,25^ and sophisticated NMR experiments have provided energetic and kinetic information on nascent chain folding. Single-molecule force spectroscopy with optical tweezers^15,26,27^ have successfully been used to control the folding and unfolding of nascent proteins at the single-molecule level, revealing energetic and kinetic aspects of co-translational folding. However, these techniques can only be applied under *in vitro* conditions that cannot recapitulate the networks of interactions that nascent chains are a part of in the cytosol. Ribosome profiling experiments, combined with immunoprecipitation (IP)^19,28–30^, have been used to assess nascent chain folding *in vivo*, but are limited by the availability and specificity of antibodies for IP. New experimental approaches are needed to define co-translational folding pathways in the context of the relevant cellular environment.

In-cell NMR^31–33^ and FRET measurements using genetically encoded fluorophores, combined with temperature jump protocols^34^ can be used to determine protein folding and stability in live cells. However, these approaches are technically very demanding, and it has not yet been possible to apply them to studies of nascent polypeptide folding on the ribosome.

We have previously developed a reporter assay for co-translational folding that exploits reversible translation arrest by the SecM17 arrest peptide (AP)^35,36^. Co-translational folding events release translation arrest, resulting in reporter expression^37^. We applied this approach to define the folding pathway of a nascent protein domain in live cells^38^, highlighting the potential of the technique for scalable, mechanistic studies of co-translational folding. In this work, we developed a high-throughput reporter assay based on folding-induced arrest release that we term “AP profiling”. We find that the homologous EF-G and EF-Tu from *E. coli* follow different co-translational folding pathways. Consistent with previous studies^38^, the G-domain from EF-G does not form a stable co-translational folding intermediate. In contrast, the homologous domain from EF-Tu exhibits a folding intermediate at a position that coincides with the insertion of the functionally important G’ subdomain in EF-G. The nascent chain-binding chaperones trigger factor (TF) and DnaK both destabilize this intermediate, suggesting that the stability of folding intermediates is modulated by both sequence context and molecular interactions. Collectively, our results demonstrate the utility of mapping co-translational folding events with codon-resolution at high throughput.

## Results

### AP profiling quantifies co-translational folding of EF-G at codon resolution

We have previously demonstrated that nascent chain folding generates forces of sufficient magnitude to overcome AP-induced translation arrest^37^. Coupling arrest release to reporter expression provides an approach for sensitive and quantitative detection of nascent chain folding in cells^37–39^. These assays required the generation and separate expression of individual clones covering the open reading frame (ORF) of the candidate protein to be investigated, limiting the throughput of the approach. To overcome this limitation, we developed a scalable version of the AP reporter assay that we termed “AP profiling”.

We generated plasmids that encode two fluorescent proteins. An mCherry^40,41^ ORF is expressed under the control of an arabinose-inducible promoter (P*_araBAD_*)^42^ and serves as an internal control for plasmid copy number and transcription levels. A separate ORF, also under P*_araBAD_* control, encodes the candidate protein fused to the SecM AP^12^ followed by GFP^43^. Candidate folding accelerates arrest release, generating more GFP and, thus, a higher ratio of the GFP and mCherry fluorescence signals (Figure 1A). This dual reporter system enables ratiometric fluorescence measurements, thus providing a robust and sensitive measurement of nascent chain folding.

**Figure 1.**
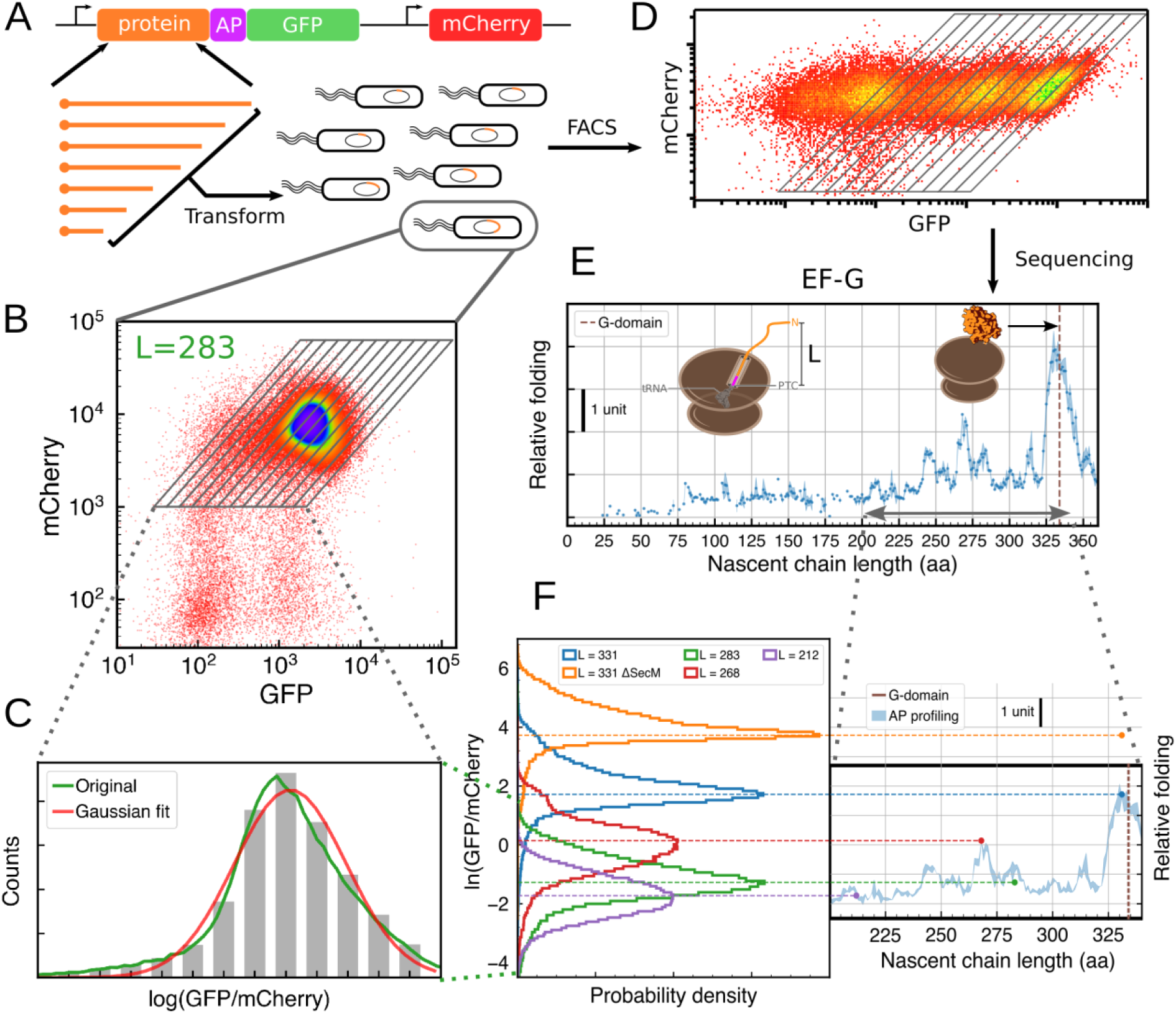
AP profiling workflow resolves co-translational folding with high throughput. (**A**) Truncation library of the candidate protein is fused upstream of arrest-peptide and GFP before being transformed into E. coli cells. (**B**) Cells carrying the same truncation fragment (L = 283 aa) display a uni-modal distribution across fluorescence levels. (**C**) The fluorescence-gated population closely follows a normal distribution. (**D**) Cells from the truncation library are sorted based on fluorescence ratio. (**E**) Plasmids in the sorted samples were extracted and sequenced to establish relative abundances of constructs across fluorescence levels before assessing folding along the ORF. (**F**) Logarithmic values of the fluorescence ratios from cytometry are consistent with the AP profiling results (200 ≤ L ≤ 240). *Δ*SecM carries P17A mutation that abolishes translation arrest and results in high GFP signals. The shaded area in AP profiling represents mean ± std from 3 repeat experiments.

To validate our approach, we first analyzed the G-domain of EF-G, referred to here as G_EF-G_. The folding of this domain has previously been investigated *in vitro*^15,26,44^ and *in vivo*^38^. We expressed AP constructs containing defined lengths of the EF-G ORF in *E. coli* cells and assessed GFP and mCherry fluorescence signals of individual cells using flow cytometry. For initial characterization, we chose lengths that are expected to yield either high (L = 331 aa) or intermediate to low (L = 212, 268 or 283 aa) GFP/mCherry intensity ratios. For all individual constructs, the logarithm of the measured fluorescence intensity ratio, log(I_GFP_/I_mCherry_), exhibited normally distributed values (Figure 1 B, C, F). A construct length of L = 331 aa results in a high GFP/mCherry ratio. To test whether this ratio is still within the dynamic range of our assay, we created a version of this construct with a defective SecM AP^35^. Abolishing arrest resulted in a further increase in GFP fluorescence, demonstrating that even the high signal from the L = 331 aa construct is still within the dynamic range of our reporter assay (Figure 1F). Taken together, our results demonstrate that the reporter construct covers a wide dynamic range, and that the log(I_GFP_/I_mCherry_) values for each individual construct are well described by a normal distribution.

The difference in the GFP/mCherry ratio among the individual constructs can be attributed to difference in the folding and stability of individual nascent chains fused to the arrest peptide, as previously described by us^37,38^ and others^25,45,46^. We thus used the mean of the log(I_GFP_/I_mCherry_) values^47^ as a measure of folding-induced arrest release, yield values of -1.73, 0.14, -1.28, 1.72 for the tested nascent chain lengths of 212, 268, 283 and 311 aa. The results described above suggest that folding scores of large numbers of constructs can be determined simultaneously when flow cytometry is combined with cell sorting. Identifying constructs in each gate using a “sort-seq” scheme^47,48^ should enable reconstruction of the normal distribution of the log(I_GFP_/I_mCherry_) values, thus making it possible to assign folding scores at high throughput.

A previous study indicated that G_EF-G_ exhibits a single major folding event around the nascent chain length of 332 amino acids (aa)^38^, using individually cloned constructs. Here, we generated G_EF-G_ truncations by time-dependent exonuclease digestion^49–51^, providing complete coverage of the candidate coding sequence without the requirement for generating individual constructs (Figure 1A). We transformed a library of G_EF-G_ truncations in our dual reporter AP construct into wild-type *E. coli* cells. After expression, we subjected the cultures to fluorescence-activated cell sorting (FACS), which revealed a distribution of GFP intensities over three orders of magnitude, whereas the mCherry signal lies within one order of magnitude (Figure 1D). Employing a robust bacteria sorting strategy (Supp. Figure 1), we sorted cells into pools of similar log(I_GFP_/I_mCherry_) values, before analyzing the plasmid DNA from individual pools with a custom deep-sequencing workflow. The relative abundance of individual library members across all sorting gates was used to calculate a “folding score” based on the mean of their distribution (see Methods). The folding scores determined in this way exhibited a single major peak at a nascent chain length of 331 aa (Figure 1E), in excellent agreement with previous findings using a different reporter system^38^. Folding scores calculated from cytometry of individual constructs matched those determined by FACS, validating our sorting and sequencing workflow (Figure 1F). Taken together, our novel approach of combining folding-induced AP release, cell sorting and deep sequencing described here detects co-translational folding events at high throughput in live cells. We term this method “AP profiling”.

Earlier work did not detect any major folding events prior to full extrusion of the G_EF-G_ from the ribosome^38^. AP profiling reproduced the main folding event for the full G_EF-G_, but also detected substantial signals at chain lengths above 230 aa (Figure 1E). Part of this apparent discrepancy results from AP profiling reporting folding signals on a logarithmic scale, leading to low-amplitude signals appearing more prominently. Linearly scaled I_GFP_/I_mCherry_ closely resemble the folding profile reported previously using a luminescent reporter^38^ (Supp. Figure 2). The improved sensitivity in AP profiling facilitates detection of minor folding signals. Single-molecule force spectroscopy experiments with optical tweezers indicated that the G_EF-G_ does not form stable tertiary structure before extrusion of the complete domain^38^. As such, the minor folding peaks at lengths of 245, 270 and 280 aa were unexpected.

To test whether AP profiling reveals previously undetected tertiary structure formation, we generated a destabilized version of G_EF-G_. We replaced all 8 phenylalanine residues with alanine residues (F/A mutant) (Figure 2C, red residues). Replacing the bulky hydrophobic side chain of phenylalanine with the much smaller one of alanine disrupts packing of the hydrophobic core of the domain, which greatly destabilizes tertiary structure^52^. The main folding peak of the F/A mutant G_EF-G_ was significantly reduced compared to that of the wild-type domain, whereas the minor folding peaks remained unchanged (Figure 2A). The difference between the signals (“ΔFolding”, Figure 2A) thus remains close to zero throughout most of the ORF and only rises near the main folding peak at L = 331 aa. These observations suggest that processes other than tertiary structure formation reduce arrest stability, perhaps including the formation of short-range interactions or secondary structures. However, stably folded tertiary structures still elicit the strongest arrest release. In line with expectations, single-molecule experiments with optical tweezers did not detect any stable tertiary structure for the mutant G_EF-G_ (Figure 2B). Thus, control experiments with a folding-defective mutant help to interpret AP profiling data and eliminate non-random fluctuations that are not related to tertiary structure formation.

**Figure 2.**
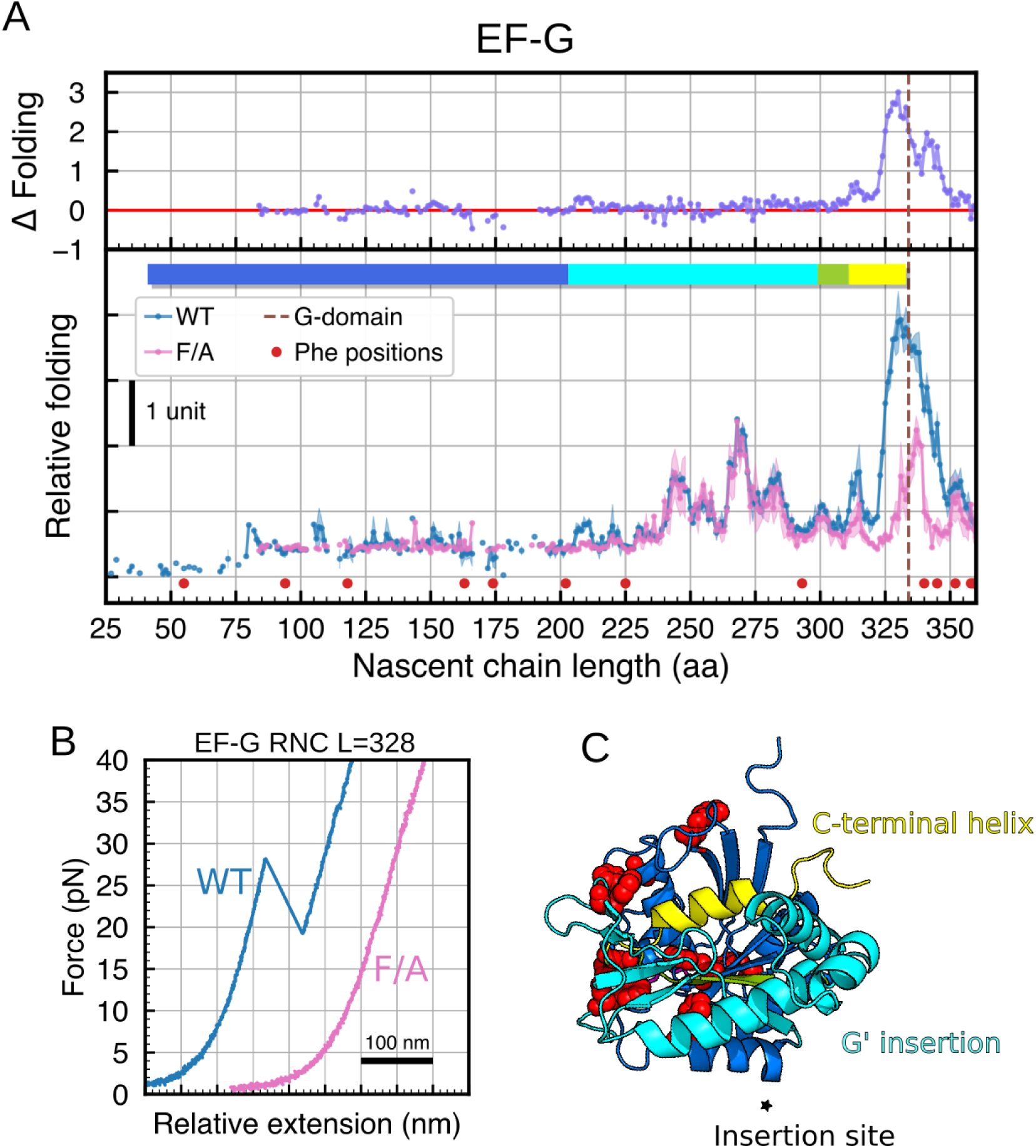
AP profiling resolves G_EF-G_ folding at codon resolution. (**A**) AP profiling of wild-type (WT, n=3) and mutant (F/A, n=2) EF-G. The dots are average values and the shaded areas are standard deviations from repeat experiments. All phenylalanine residues were mutated to alanine in the F/A mutant. Red dots label the nascent chain lengths when the mutated residues are incorporated into the peptide. Blue, cyan and yellow bars indicate when the structural elements (matching panel **C**) are extruded from the ribosome, assuming 40 aa residing in the ribosome exit tunnel. Folding differences are calculated by subtracting mean values. (**B**) Single-molecule force spectroscopy of the EF-G ribosome-nascent chain complex (RNC) at L = 328 aa. The WT EF-G shows a clear unfolding transition when applied force while the F/A EF-G does not have a stable structure. (**C**) Molecular structure of G_EF-G_ (adapted from PDB entry 7PJV), residues colored in red are phenylalanines. Green indicates the position of a connecting strand just before the C-terminal helix.

### Homologous EF-Tu has an on-pathway folding intermediate

We previously conjectured that the particular folding order of the G_EF-G_ is due to the insertion of the G’ subdomain that wraps around the C-terminus of the domain^38^. EF-G belongs to a superfamily of translational GTPases (trGTPases) that share a conserved domain topology^53–55^. Elongation factor Tu (EF-Tu) represents another highly abundant and essential trGTPase^56,57^ that shares structural homology with EF-G. However, the G-domain of EF-Tu (G_EF-Tu_) does not have the G’ insertion (Supp. Figure 3). To investigate the folding of *E. coli* G_EF-Tu_, we constructed AP profiling libraries of the wild-type (WT) sequence as well as a folding-deficient mutant (F/A) (Figure 3C). Both the wild-type and the mutant candidate showed multiple folding peaks along the coding sequence (Figure 3A). Repeated experiments in EF-Tu consistently missed coverage in certain parts of the open reading frame, suggesting growth bias or toxicity from some EF-Tu fragments. Nevertheless, we obtained signals for most positions within the ORF. The mutant EF-Tu sequence exhibited lower folding scores than wild-type in several regions, resulting in positive differential signals (Figure 3A, “ΔFolding”), in contrast to what is observed for the EF-G sequence.

**Figure 3.**
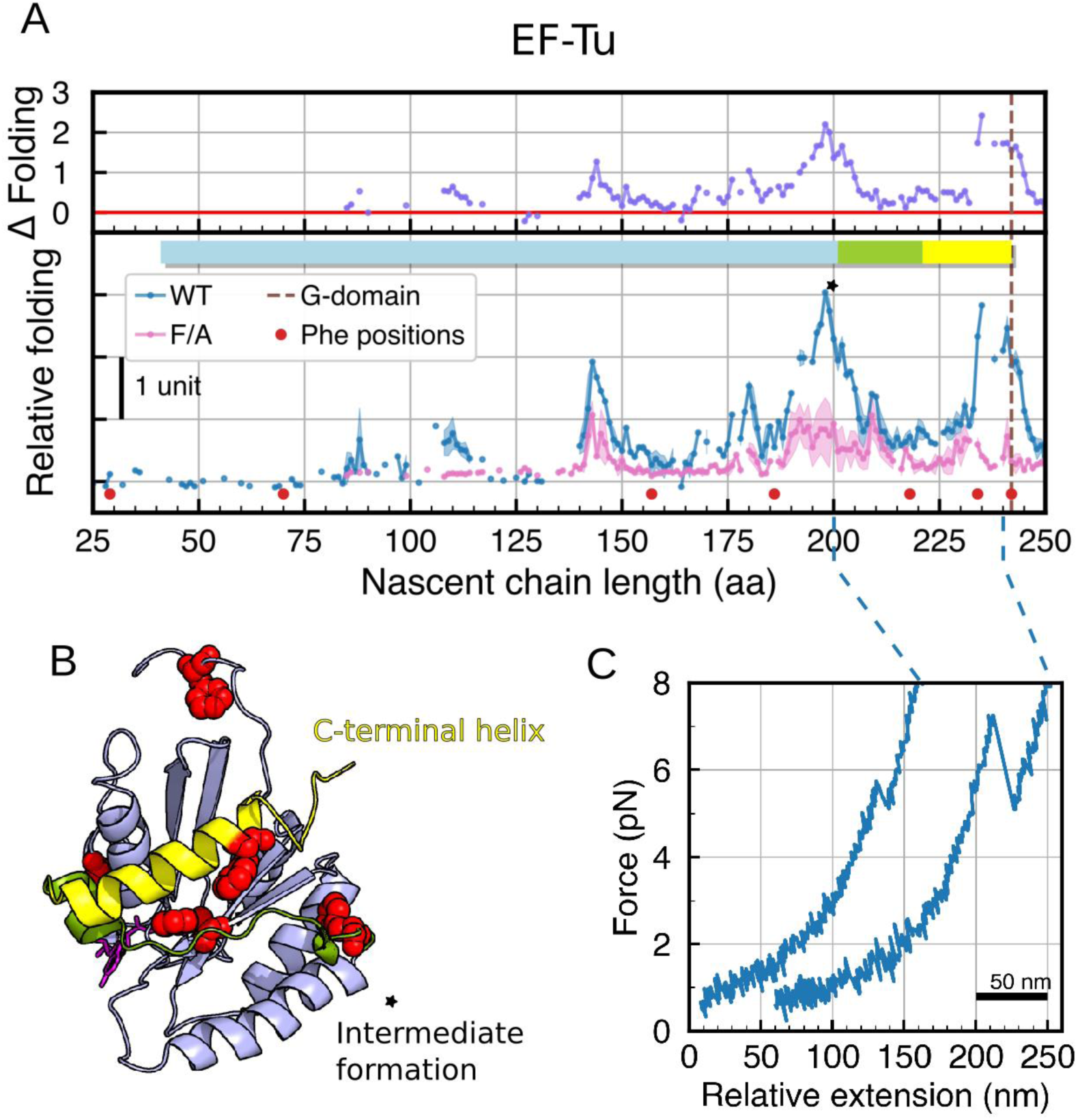
EF-Tu has a stable folding intermediate before the entire domain is synthesized. (**A**) AP profiling results of the wild-type and F/A EF-Tu proteins. Shaded areas represent standard deviations from repeat experiments (n=2). Red dots match nascent chain lengths when the mutations are incorporated into the peptide. Folding differences are calculated by subtracting mean values. (**B**) Molecular structure of EF-Tu (adapted from PDB entry 5UYL), residues colored in red are phenylalanines. (**C**) Single-molecule force spectroscopy of EF-Tu RNCs at L = 200 and 240 aa. Representative traces were plotted at 125 Hz to highlight unfolding transitions.

Near the position of full domain extrusion, from the ribosome (L = 242 aa), a strong signal was observed for wild-type G_EF-Tu_. The F/A mutant exhibited significantly reduced signal at this length, indicating stable tertiary structure formation at this position. Additionally, strong signals were observed for the wild-type but not the F/A construct at L = 143 aa and L = 198 aa. At L = 143 aa, the molecular structure indicates that the second to last strand of the main sheet is extruded, corresponding to a medium level folding peak. The nascent chain at L = 198 aa corresponds to the extrusion of the second to last helix.

To confirm structure formation at positions indicated by the AP profiling results, we carried out single-molecule force spectroscopy with nascent EF-Tu polypeptides (Figure 3B). At a nascent chain length of L = 240 aa, the G_EF-Tu_ is fully extruded from the ribosome and should fold into its native structure. Consistent with this expectation, we observed a cooperative unfolding transition of 69.6 nm contour length for ribosome-nascent chain complexes with a 240-amino acid long EF-Tu nascent chain. At a nascent chain length of L = 200 aa, a smaller unfolding transition of 27.3 nm contour length was observed, indicating the formation of tertiary structure at. Our optical tweezers measurements thus confirm the formation of tertiary structure within the partially extruded G_EF-Tu_ that is observed in our AP profiling experiments.

A previous study found that the G_EF-G_ folds without populating a co-translational folding intermediate before the full domain is extruded from the ribosome^38^. This observation was rationalized by noting that the G’ insertion in G_EF-G_ engulfs the helix at the extreme C-terminus of the domain and might require the C-terminal helix to fold before the G’ insertion^38^. This interpretation is supported by our findings presented here. G_EF-Tu_ lacks the G’-insertion and does not appear to be constrained in forming co-translational folding intermediates. The two highly homologous proteins thus fold co-translationally along distinct pathways that may be constrained by their respective topologies.

### Molecular chaperones disrupt folding intermediates in trGTPases without G’ insertion

Molecular chaperones are essential for efficient cellular protein folding^58^. A number of (typically small) proteins refold efficiently *in vitro* without assistance^1^. However, genetic ablation of chaperones results in large-scale misfolding and aggregation^19^, underscoring their importance *in vivo*. Despite their uncontested importance, it is not clear how chaperones act to promote efficient folding. It is conceivable that long stretches of unstructured polypeptide, such as the incomplete G_EF-G_, are stabilized by chaperones to prevent aggregation and misfolding. In addition, chaperones might help to prevent the population of kinetically trapped off-pathway folding intermediates^59,60^ that reduce folding cooperativity and slow down overall folding kinetics^61^. With its ability to detect co-translational structure formation, AP profiling might offer an opportunity for defining chaperone contributions to nascent protein folding *in vivo*.

In *E. coli* and most other bacteria, trigger factor (TF) and DnaK are the main chaperones that interact with the growing nascent polypeptides^4,58^. The ATP-independent TF binds to a specific site on the ribosome adjacent to the tunnel exit and interacts with the majority of nascent chains^28^. DnaK, the bacterial Hsp70 homolog, requires ATP and the co-chaperone DnaJ for regulated cycles of substrate binding and release that facilitate nascent protein folding^62^. Genetic studies suggest that these two chaperones are at least in part functionally redundant, with overlapping substrate pools^19,63,64^. Individual deletions are tolerated, whereas combined ablation of both chaperones results in severe temperature sensitivity and growth defects^65^. However, whether TF and DnaK employ similar or distinct mechanisms to chaperone nascent chain folding remains unknown.

To detect specific chaperone effects along the lengths of G_EF-G_ and G_EF-Tu_, we carried out AP profiling experiments in TF and DnaKJ deletion strains (*Δtig* and *ΔdnaKJ*, respectively). Ablating TF or DnaKJ did not significantly affect folding signals of G_EF-G_ (Figure 4A). The magnitude of G-domain folding at full extrusion (L = 331 aa) remained unchanged and the measurements showed only relatively minor folding score differences further upstream in the open reading frame. Small but noticeable increases in folding scores are observed in the region of the G’ insertion when we removed TF (L

**Figure 4.**
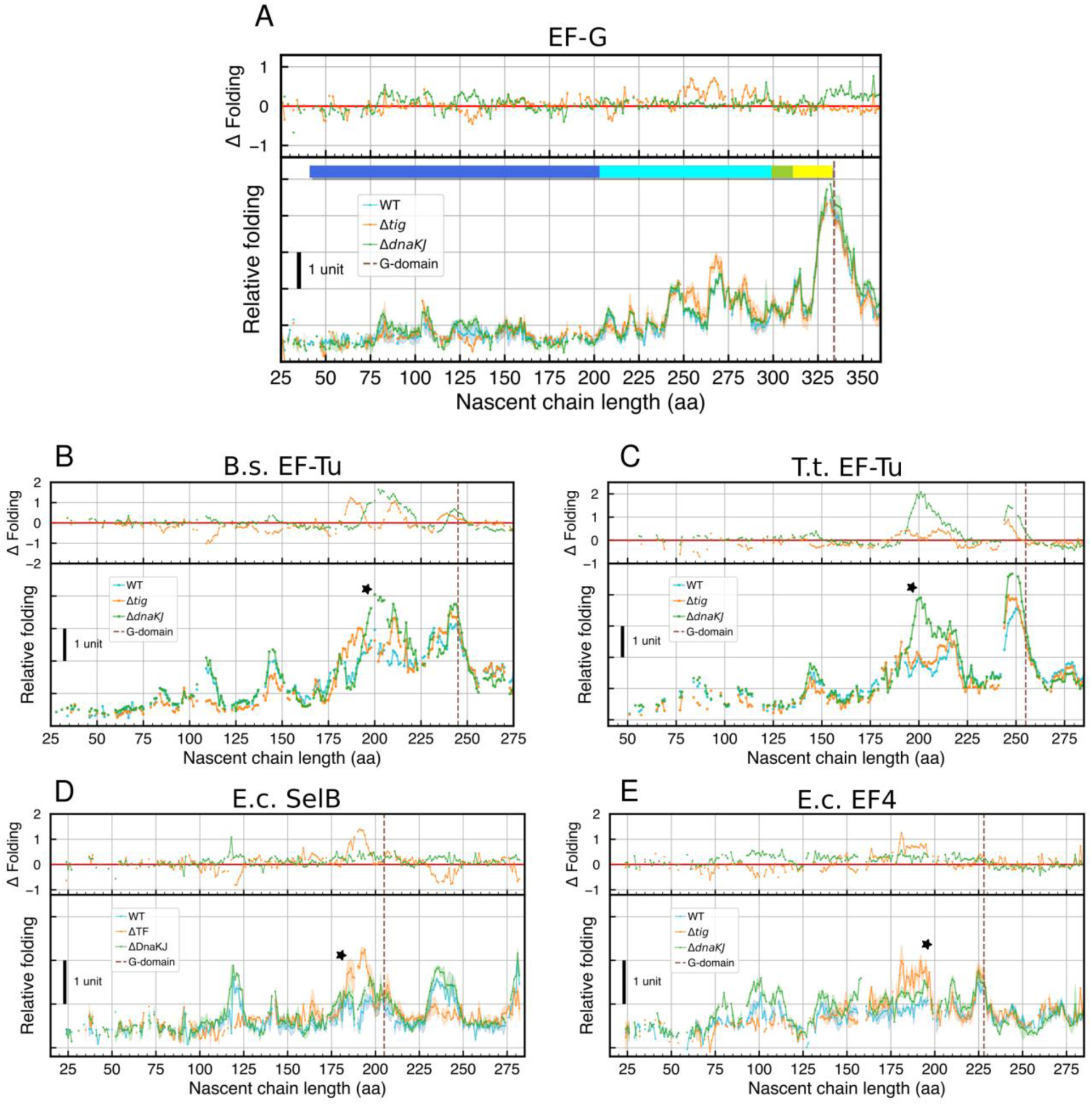
Chaperone interactions in EF-G homologs revealed by deletion strains. (**A**) AP profiling results (n=3) of G-domain from EF-G showed minimal chaperone effects on G-domain folding. (**B-E**) Chaperone effects in folding of G-domain homologs. EF-Tu from B. subtilis (strain 168) and T. thermophilus (HB8 strain) (n=1) showed clear chaperone interaction effects at the intermediate sites. Translational GTPases without the G’ insertion from E. coli, SelB and EF4 (n=3), also showed significant chaperone deletion effects at the similar structural position.

= 255, 270, 280 aa). Interestingly, these elevated peaks were not disrupted by extensive mutagenesis in EF-G (Figure 2A, F/A mutant) and thus are likely representing secondary structure formation. Our results indicate that TF binding at these positions reduces the formation of these structures.

While chaperone deletions caused only moderate effects on G_EF-G_ folding, pronounced changes were observed for G_EF-Tu_ (Figure 4 B and C). However, poor coverage makes a detailed evaluation difficult (Figure 3A). To overcome this challenge afflicting *E. coli* EF-Tu, we carried out AP profiling experiments with EF-Tu orthologs from *Bacillus subtilis* and *Thermus thermophilus*, which are highly similar to the *E. coli* EF-Tu (72% and 74% sequence identity, respectively). Experiments with *B. subtilis* and *T. thermophilus* EF-Tu yielded good coverage and, importantly, exhibited similar features that were observed for the *E. coli* protein: a peak around L = 200 aa, corresponding to a co-translational folding intermediate, and a peak around L = 250 aa, corresponding to the fully extruded G_EF-Tu_.

In the *Δtig* and *ΔdnaKJ* strains, G_EF-Tu_ folding signals were elevated at several positions compared to wild-type cells. Intriguingly, the intermediate formation site (Figure 4 B and C; L = 200 aa, star) exhibited drastically increased folding signals when either TF or DnaK was deleted, suggesting that cellular TF and DnaK destabilize the intermediate. Competition with folding or misfolding is a well-documented feature of chaperone function, although a more active role in folding has also been suggested^66–68^. At some positions in the EF-Tu coding sequence, the folding signals in the *Δtig* strain were lower than in wild-type cells (Figure 4B, L = 108 aa and L = 145 aa), which might suggest the formation of local structures protected by TF^69^ or elevated arrest release induced by direct TF binding.

In addition to EF-Tu, other homologous trGTPases contain G-domains without G’ insertion. We subjected two of these homologs, SelB^70^ and EF4^71,72^ to AP profiling experiments. Both SelB and EF4 showed G-domain folding peaks (Figure 4D, L = 205 aa; Figure 4E, L = 228 aa). Both proteins also exhibited an intermediate folding region, which was elevated in chaperone deletion strains (Figure 4 D and E, star). This pattern of intermediate formation and chaperone effects in the P-loop GTPases suggests that this folding intermediate is a major substrate of the chaperone TF and DnaK and perturbing the intermediate might be beneficial for productive folding of the domain and the full protein.

### AP profiling resolves full-length domain-wise co-translational folding in vivo

Previous folding measurements using arrest peptides employed *in vitro* translation systems^25,45,46^. In contrast to full cellular translation, *in vitro* translation systems synthesize only relatively short polypeptides efficiently. For longer translation products, accumulation of incomplete products is commonly observed^45^. Application of arrest peptide-based folding measurements *in vitro* is thus limited to small proteins or domains. Since AP profiling is an *in vivo* approach, it does not suffer from this limitation and should be capable of resolving co-translational folding of long proteins, such as the 704-amino acid full-length EF-G (Figure 5A).

**Figure 5.**
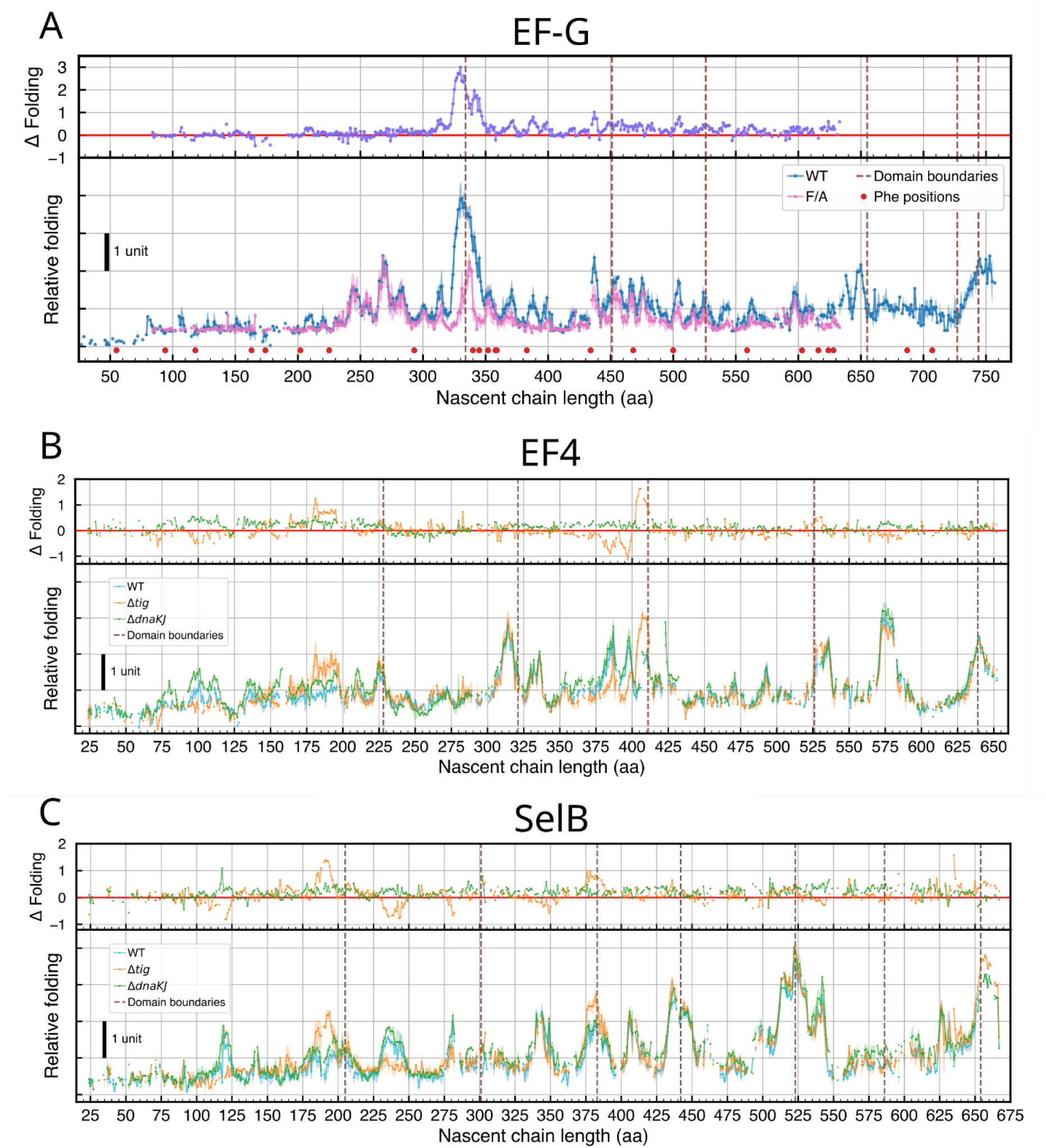
AP profiling resolves full-length domain-wise co-translational folding in vivo. (**A**) G-domain and domain II in EF-G exhibited domain-wise folding while Domain III does not show clear folding signal, consistent with previous observation (Liu et al., 2019a). Full-length protein folding can be observed in the end. The original protein open reading frame (704 aa for EF-G) is extended with glycine-serine linker (65 aa) in order to observe full-length protein extrusion from the ribosome. We were only able to retrieve the EF-G F/A truncation library until the first 610 residues in the E.coli TOP10 strain. WT (n=3) and F/A (n=2) were visualized in the same way as in Figure 2. (**B-C**) Domain-wise folding events of full-length EF4 and SelB (n=3) can be resolved in AP profiling in the MC4100 strains, highlighting the specific chaperone dependencies.

Single-molecule studies revealed that the five-domain EF-G folds according to a prescribed order^15,44^. The two N-terminal domains (G and II) domains fold co-translationally as they are being extruded from the ribosome, while the C-terminal domains III, IV and V fold only after the entire protein is synthesized^44^. AP profiling of WT and F/A variants of EF-G provided a full-length profile of the 703-residue EF-G folding (Figure 5A). Domain II folding signals did not rise to the same magnitude as those of the G_EF-G_, but the several smaller peaks are detected before and near domain II boundary (L = 451 aa), suggesting that domain II folds in multiple steps at chains lengths of L = 370 aa and L = 385 aa. Domain III did not show significant folding peaks near the domain III boundary (L = 526 aa), consistent with previous finding that domain III folds post-translationally, because it requires interactions with its C-terminal neighbors for tertiary structure stability^44^. We were unable to acquire data for the F/A mutant at chain lengths above L = 630 aa because of cloning difficulties. Full-length AP profiling results of SelB and EF4 were obtained in chaperone deletion strains. We observed domain-wise co-translational folding events from all five domains in EF4 at individual domain boundaries (Figure 5B). At the C-terminus, one additional folding peak at L = 575 aa maps to a beta-hairpin formation, which is proposed to induce strong arrest release^73^. Interestingly, the absence of TF decreased folding signal at positions L = 385 aa and L = 395 aa, but generated a much larger folding peak at L = 408 aa. One possible explanation is that TF stabilizes smaller secondary structures which results in less cooperative domain folding at L = 411 aa and thus lower folding signals. In contrast to EF4, SelB did not exhibit folding at all domain boundaries (Figure 5C). Notably, we did not detect significant folding signals for domain II (L = 300 aa) and domain VI (L = 585 aa), highlighting that domains do not always fold co-translationally, as has previously been observed in single-molecule measurements^44^. Our results demonstrate that AP profiling can resolve complex co-translational folding pathways of large multi-domain proteins.

## Discussion

Arrest peptides have recently found widespread use as sensors of co-translational folding^22,22,25,38,39,45,46,52,74–76^. Here, we have combined this approach with reporter expression, ratiometric flow cytometry, cell sorting and deep sequencing to build a sensitive, high-throughput approach for mapping co-translational folding with codon resolution. We termed this method Arrest Peptide profiling (AP profiling).

Here we have utilized AP profiling to investigate the co-translational folding of homologous trGTPases in living cells. Following co-translational folding in cells with AP profiling enables studies of large, multi-domain proteins that are inaccessible by *in vitro* translation systems, which synthesize only relatively small proteins efficiently^46,76^. In addition, cellular measurements, combined with genetic manipulation, capture nascent chain-chaperone interactions, and reveal their effects on co-translational folding.

Our AP profiling measurements readily reproduced the earlier observation that G_EF-G_ does not form a stable folding intermediate before the full-length domain is extruded from the ribosome^38^. Previous approaches required cloning of discrete constructs that were expressed individually. In contrast, the AP profiling approach developed here allows studies in a library format. Moreover, AP profiling has higher sensitivity than previous methodologies, resulting in detection of previously unresolved signals (Figure 2A, cyan region). Using folding-defective mutants, we show that specific signals do not reflect tertiary structure formation. Rather, they can be attributed to secondary structure formation, which can take place in^77^ and even be stabilized by^11^ the ribosome exit tunnel. The latter observation might also explain why introduction of several glycine substitutions, typically regarded as helix breaking substitutions^78^, does not disrupt bona-fide helix formation (Supp. Figure 4). Regardless of their origin, we were able to resolve previously undetected signals in live cells, highlighting the sensitivity and large dynamic range of our AP profiling approach.

The sensitivity and throughput of our newly developed approach enabled us to follow co-translational folding of EF-G homologs, comparing the folding pathways of related proteins. The G_EF-Tu_ lacks an insertion (G’ subdomain) present in EF-G and exhibited folding through co-translational intermediates (Figure 3A). Significant folding peaks earlier than the full domain extrusion likely reflect P-loop *β*-sheet formation (L = 143 aa) as well as folding of an intermediate (L = 198 aa) at the position of G’ domain insertion. G-domains from the SelB and EF4 proteins, both of which lack the G’ insertion, showed clear folding at the same positions (Figure 4 D and E).

The G’ insertion in EF-G is functionally important and interacts with ribosomal proteins L7 and L12^79,80^. The trGTPases studied here that lack the G’ insertion (EF-Tu, SelB and EF4) all exhibit a strong co-translational folding intermediate. The position of the intermediate coincides with the site of G’ insertion. The appearance of the G’ insertion during evolution might have necessitated the elimination of the folding intermediate for topological reasons^38^, thus profoundly affecting the co-translational folding pathways of upstream sequences in EF-G (and potentially other G’-containing trGTPases).

To explore how chaperones guide co-translational folding, we carried out AP profiling experiments in *E. coli* strains harboring deletions in nascent chain-binding chaperones. Removal of the nascent-chain binding TF or DnaK chaperones resulted in minimal changes of folding intensities across the G_EF-G_ (Figure 4A), suggesting that its folding is chaperone-independent, but also that these chaperones are not required to maintain the domain in an unfolded, folding-competent state during the relatively long period of chain elongation before full extrusion and folding. However, chaperone deletions elevated folding intensities for the full domain as well as the previously detected intermediate in EF-Tu proteins from several bacterial species (Figure 4 B and C). TF and DnaK thus destabilize the intermediate formed in nascent EF-Tu.

*In vivo* measurements provided full coverage of large multi-domain proteins such as EF-G, EF4 and SelB in AP profiling (Figure 5). The folding signals from EF-G pointed to co-translational folding of the N-terminal G and II domains and post-translational folding of the domain III, consistent with the folding pathway determined by single-molecule optical tweezers^15,44^. EF4 exhibited co-translational folding from all five domains at individual domain boundaries with TF interaction detected at domain III (Figure 5B). In contrast, SelB did not exhibit domain-wise folding at domain II and VI (Figure 5C). The diverging folding pathways from homologous proteins highlighted the nuances of multi-domain protein folding and the need for more thorough and systematic examination of these prevalent large proteins.

From the candidate proteins we examined in AP profiling, DnaK deletion only resulted in higher folding signals (Figure 4) while TF deletion showed position- and context-specific effects with evidence of both higher and lower folding signals. Rescue from kinetic traps by DnaK^62,81^ and by TF^18,82^ has been reported to facilitate full protein folding for complex folders. Supplementing an *in vitro* translation system with TF was also shown to reduce folding-induced arrest release of DHFR^83^, suggesting TF disrupts folded structures. Another *E. coli* chaperone, SecB, also exhibited strong anti-folding activity^84^. Our AP profiling results from EF-G and EF-Tu obtained similar observations where TF destabilizes folded structures (Figure 4). TF has been suggested to exhibit diverse modes of action. Its substrate binding cavity^69,85,86^ might allow for limited structure formation and thus cause elevated folding signals (Figure 4B, EF-Tu L = 110 aa; Figure 4D, SelB L = 120 aa). Smaller folding events are less likely to be detected distant from the ribosome even when they are induced by DnaK interactions as folding-induced arrest release requires proximity to the ribosome surface.

One limitation of the cellular AP profiling experiments that we encountered are growth defects caused by specific constructs. Reliable folding detection requires comparable cell viability and growth rates for all truncation fragments within the library. Expressing arrest peptides generates stalled ribosomes which can interfere with growth^87^. In addition, fragments of some proteins can cause cell toxicity^88^. In our experiments, we repeatedly observed loss of coverage in some regions of EF-Tu (Figure 3A). Particularly, we were unable to obtain truncation libraries of specific candidate proteins in our *E. coli* strains (RF3 and *E. coli* EF-Tu, etc.). However, the library viability seems strain-specific, for example, *E. coli* EF-Tu truncation library is not viable in MC4100 strains, but is viable in the TOP10 *E. coli* strain (Figure 3A). The folding scores of the same protein are also consistent across different wild-type strains (Supp. Figure 5). This suggests that strains for AP profiling experiments can be tailored to specific candidate proteins for optimal coverage.

In summary, we developed a powerful tool, AP profiling, for following co-translational folding in live bacterial cells with high throughput. We employed the approach to study homologous trGTPases, including the abundant EF-G and EF-Tu proteins. Our experiments revealed a divergence in co-translational folding pathways that results from a structural insertion. Chaperone interactions caused complementary effects on disrupting folding intermediates. Our approach can readily be extended to study other aspects of nascent chain biology, such as membrane insertion^89,90^. The unparalleled throughput and sensitivity make AP profiling ideally suited for *in vivo* studies of protein folding and membrane insertion and to dissect contributions from chaperones and other cellular machinery. Collectively, our advances provide much-needed tools for mechanistic investigations of protein biogenesis.

## Methods and Materials

All commercially available enzymes were purchased from NEB unless stated otherwise. PCR reactions were carried out with Phusion high-fidelity DNA polymerase (Thermo Scientific, F530S). Chemicals were purchased from Sigma-Aldrich unless stated otherwise. DNA Tini spin column (Enzymax LLC, EZC106N) was used for DNA purification protocols. GeneJET plasmid miniprep kit (Thermo Scientific, K0503) was used for all plasmid miniprep steps. GeneJET PCR purification kit (Thermo Scientific, K0701) was used for all dsDNA purification, unless stated otherwise. The binding buffer from K0701 was tested and used for quenching enzymatic reactions in AP profiling at twice the reaction volume. *E. coli* TOP10 cells was from a commercial source (Invitrogen, C404010). MC4100 strains were from a previous study^65^. Double stranded DNA fragments (gBlock) and sequencing adaptors are synthesized by IDT (Integrated DNA Technologies). Detailed information about candidate proteins used in this study is listed in **Table 1**.

**Table 1.** Detailed information on protein candidates used in this study. Proteins E. coli are cloned from K-12 strain. Proteins from T. thermophilus are cloned from HB8 strain. Proteins from B. subtilis are cloned from strain 168.

| Uniprot | Name | Species | Length | Structure | Domain boundaries |
| --- | --- | --- | --- | --- | --- |
| P0A6M8 | EF-G | *E. coli* | 704 | 7PJV | 294,411,486,615,687,704 |
| P0CE47 | EF-Tu | *E. coli* | 394 | 5UYL | 202, 296, 395 |
| P14081 | SelB | *E. coli* | 614 | 5LZD | 165,261,343,402,483,546,614 |
| P60785 | EF4 | *E. coli* | 599 | 3CB4 | 188,281,371,486,599 |
| P60338 | EF-Tu | *T. thermophilus* | 406 | 1EXM | 215 (G-domain) |
| P33166 | EF-Tu | *B. subtilis* | 396 | AF-P33166-F1 | 205 (G-domain) |
| P13551 | EF-G | *T. thermophilus* | 691 | 4M1K | 284 (G-domain) |
| P80868 | EF-G | *B. subtilis* | 692 | 5VH6 | 282 (G-domain) |

### Truncation library construction

Truncation library is built by time-dependent exonuclease digestion, adapted from protocols described previously^49–51^. Purified plasmids were digested stepwise with SacI (NEB, R3156S) to prevent exoIII digestion on the arrest peptide side and then with AvrII (NEB, R0174S) to allow for exoIII digestion towards the candidate protein. Linearized dsDNA was purified and quantified by Qubit (1X dsDNA high sensitivity assay kit, Invitrogen, Q33231). Incremental digestion by exoIII (Promega, M1811) was performed by diluting the linearized plasmids to a final concentration of 33 ng/ul before adding the exoIII amount to final 0.1U per nanogram of DNA (Qubit reading). The reaction mixture was left in a heatblock at 22 °C for 5 min before sampling every 30 s into a chilled quenching buffer (binding buffer from Thermo Scientific, K0701). To blunt the digested DNA, mung bean nuclease (NEB, M0250S) was mixed with the purified DNA at a final concentration of 1.5 U per microgram of DNA to blunt the DNA and digest the remaining RNA from miniprep without introducing non-specific cleavages in the dsDNA region. Klenow fragment (NEB, M0210S) was added to the purified DNA for end-polishing at the final concentration of 1 U per microgram of DNA. The end-polished DNA was then diluted for blunt-end intramolecular ligation by T4 DNA ligase (NEB, M0202S). Ligation reaction was carried out at 16 °C for 16-20 hr. The circularized DNA was then purified carefully by washing multiple times during a standard purification protocol and eluted to a final concentration around 40 ng/ul.

Transformation efficiency was tested by transforming 50-60 ng ligation product into TOP10 cells by electroporation. Importantly, all transformations were done with freshly cultured cells. One transformation reaction consists of 5 ml fresh culture harvested at OD_600_ around 0.4. In our workflow, harvesting the cells in cold temperature gave us more consistent transformation results, while room-temperature harvest offered comparable or sometimes higher yield as reported previously (Tu et al. 2016). Cells were cultured in LB supplemented with 1% glucose at appropriate temperatures (37 °C for TOP10, 30 °C for MC4100 strains) to recover for 2 hr at 220 rpm. Cultures were spun down and spread on agar plates supplemented with 1% glucose (to repress P_BAD_ expression^91^) and carbenicillin plus strain-specific antibiotics^65^. Plates were imaged for colony counting facilitated by openCFU^92^. Library coverage was calculated based on the potential truncation fragments (in-frame and out-of-frame constructs) and colony counts. Large-scale transformation was done for the truncation library to reach at least 1000X coverage. Colonies were then scraped and resuspended in fresh LB before depositing as glycerol stocks for long-term storage.

### Bacteria fluorescence activated cell sorting (FACS)

Glycerol stocks of the truncation library in TOP10 cells were revived for overnight cultures in LB supplemented with 1% glucose. Cells carrying the truncation library were freshly diluted into glucose-free medium and grown for at least 2 hr to OD_600_=0.2. The cultures were induced with arabinose at the final concentration of 0.05% (w/v) to express for 2 hr.

The bacteria population was diluted to PBS and sorted with a MoFlo XDP equipment (Beckman Coulter) based on GFP and mCherry signals. Based on our AP profiling workflow and a similar bacteria sorting application^93^, the first sort of the population (enrichment sort) serves to enrich the scarce high GFP population. We arbitrarily set up four gates for high mCherry events, which contain the expressing population, from low to high GFP signals. Sorting was done at around 14,000 events per second and typically proceeded for 10 min when the scarcest gate contains at least 20,000 sorted events. Sorted samples were grown in LB supplemented with 1% glucose (w/v) and carbenicillin plus strain-specific antibiotics overnight at 37 °C, 220 rpm. Plasmids and glycerol stocks were prepared from the overnight cultures.

For AP profiling in TOP10 cells, glycerol stocks of TOP10 cells were revived for fresh culture before mixing in proportions to enrich high GFP species. For AP profiling to study chaperone effects, purified plasmids were mixed in proportions to enrich high GFP species before being electroporated into MC4100 strains (PG1 for WT, PG2 for *Δ*TF and PG3 for *Δ*DnaKJ) with minimum 1000X coverages. The new mixed cultures were then sorted again (quantification sort) with a more careful gating strategy (Figure 1D), similar to previous studies^94,95^. Specifically, the log(GFP/mCherry) values linearly carve up the GFP-mCherry plot on the log scale during cytometry. The Summit (v5.5.0) software allows for manually adjusting gate boundaries according to a coordinate system. We then drew up 12 slanted (slope = 1) gates on the high mCherry population, shoulder by shoulder, with a fixed gate width. Each sorting gate now can be represented by a single log(GFP/mCherry) value. The MoFlo XDP allows for sorting 4 gates simultaneously. We ran the sorter at around 14,000 events per second and recorded the exact numbers of detected events, sorted events and the total number of events in the high-mCherry cluster (for normalization across groups during analysis). Running the sorted samples back through the cytometry resulted in reproducible signal levels, verifying singlet bacteria sorting (Supp Figure 1). Each 4-gate sorting lasts around 10 min to collect enough cells for sequencing detection. Sorted samples were grown in LB supplemented with 1% glucose (w/v) and carbenicillin plus strain-specific antibiotics overnight at appropriate temperatures (37 °C for TOP10, 30 °C for MC4100 strains), 220 rpm. Plasmids and glycerol stocks were prepared from the overnight cultures.

### High-throughput sequencing

We developed a custom end targeting high-throughput sequencing scheme, inspired by Tail-seq^96^. Plasmids extracted from truncation library constructions, transformations and sortings were used for sequencing library preparations. Briefly, the BsaI recognition site near the arrest peptide coding sequence was digested and the overhung on the truncation fragment site was used to ligate with a biotin modified P5 adaptor containing the UMI and desired index. The single stranded portion of the P5 adapter was then filled up by Klenow fragment before purification and quantification by Qubit. We subcloned the Tn5 transposase (from Addgene plasmid #60240) and purified the enzyme for DNA tagmentation^97,98^. The ME-B fragment was transposed into the DNA samples before biotin enrichment by streptavidin-coated magnetic beads (M280 beads, Invitrogen, 11205D). The beads were vigorously washed with wash buffer according to the manufacturer’s recommendations. The magnetic beads, bound with the biotin labeled DNA fragments, were directly used as template for a limited-cycle PCR (8-10 cycles) with Phusion DNA polymerase to attach the desired P7 adaptor. PCR products were purified and subject to size selection by AMPure beads (Beckman Coulter, A63880) to remove short DNA fragments (0.8X volume).

The resulting DNA architecture is compatible with Illumina sequencing platforms. The samples from this study were sequenced with HiSeq2500 (50 cycles single end) as well as NovaSeq6000 SP (100 cycles, single end) with additional index cycles to acquire UMI information.

### AP profiling data analysis

The sequencing data was processed by custom scripts chaining standard analysis tools. Briefly, we used *fastp*^99^ for quality assessment and initial filtering of raw FASTQ reads. The two index reads were combined and used together to demultiplex the entire sequencing experiment with a custom python script. UMI bases were extracted with *umi-tools*^100^ and alignment reference files were generated according to the protein candidates included in the truncation library. Short-read sequence alignment was done with *bowtie2*^101^ and formatted with *samtools*^102^. The SAM/BAM file entries were then deduplicated by *umi-tools* if the reads shared the same UMI while being mapped at the same location. The deduplicated mapping information was then compiled into read counts in positions along the protein coding sequences from the truncation library with a custom python script. The percentage of a read count in a certain position from a sort gate was then converted to a “normalized event count” for that position using the normalized sorting information.

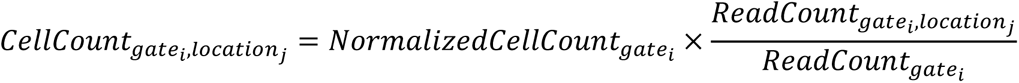

Thus, for all the mapped positions possible from the truncation library (representing all the possible truncation constructs), we acquired 12 values of “normalized event count” for each position, corresponding to 12 sorting gates. Fitting a Gaussian curve to the 12 values^47^, we acquired the peak of the curve among the 12 gates as a raw folding score. We filtered the low coverage positions and bad fits before converting the raw folding scores with the sorting gate width information to acquire the relative log(GFP/mCherry) values as adjusted folding scores.

The folding scores can be linearly adjusted and aligned across different sort experiments because of the nature of fluorescence detection and logarithmic transformation.

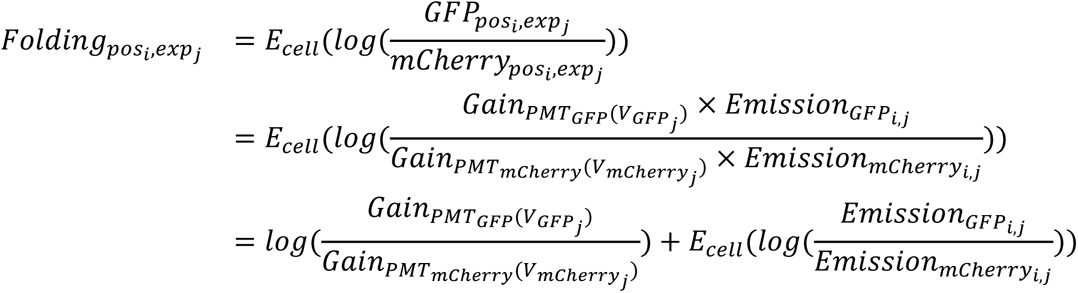

The folding score of a defined position (*pos*_*i*_) from one experiment (*exp*_*j*_) can be defined as the expected value of the log(GFP/mCherry) from the cells carrying this truncation construct. Fluorescence detection in the cytometer from the experiment can be described as the excited signals multiplied by the gain setting in the photomultiplier tube (PMT) at the detector side. The different gains of the same PMT are dependent on the voltage settings.

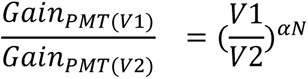

where *⍺* is a coefficient determined by the dynode material and geometry, and *N* is the number of dynodes in the PMT^103^. The gain settings in the cytometer can thus be expressed as below:

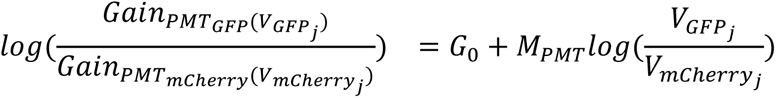

*G*_0_ is a constant independent of experimental parameters containing the base gains of the two PMT. *M*_*PMT*_ is a constant only influenced by the physical properties of the two PMT.

The fluorescence signals are proportional to the concentrations of fluorescent proteins in the cells and the inherent properties of GFP and mCherry. Assuming the same level of transcription of the two open reading frames, mCherry concentration is determined by translation initiation in the mCherry, which is constant across positions for a specific experiment, and GFP concentration is mainly determined by folding-induced arrest release, which is constant across experiments for a specific position.

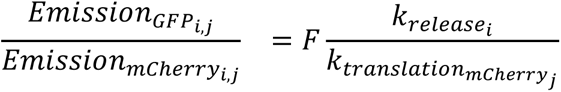

where *F* is a compound parameter containing fluorescence characteristics of GFP and mCherry, such as quantum yield. The folding score can then be expressed with experiment-specific and position-specific parameters shown below:

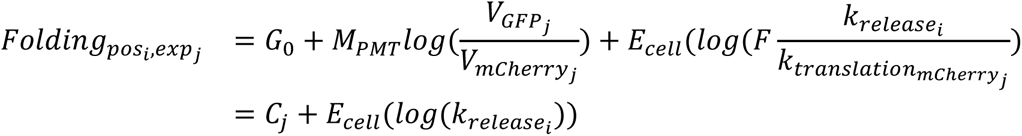

Here, *C*_*j*_ is a compound parameter containing all the experiment-specific settings, such as the PMT voltages during the sorting experiment and mCherry expression level for that experiment. The acquired folding scores from fitting Gaussian curves, *E*_*cell*_(*log*(*k*_*releasei*_)) can thus be modified linearly by adding or subtracting values to align with folding scores acquired in different experiments.

Aligned folding scores were then plotted against the nascent chain lengths of the corresponding truncation constructs (Figure 1E) for further interpretation. We used natural base for logarithmic transformation and the relative folding value of 1 between two truncation constructs means that the difference of *ln*(*GFP*/*mCherry*) values from the two constructs is 1.

### Flow cytometry of individual truncation constructs

Individual truncation constructs were generated by Gibson assembly. TOP10 *E. coli* cells were transformed with the truncation plasmids separately by heatshock. Individual colonies were used to inoculate fresh LB media supplemented with 1% glucose and carbenicillin for overnight cultures. Dilutions were made in fresh media without glucose and cultured for at least 2 hr before induction by arabinose in the same way for FACS experiments. Cells were diluted in PBS before running through an Attune NxT Flow Cytometer (Invitrogen). FCS files were analyzed by custom python scripts.

### Single-molecule force spectroscopy of RNC

Ribosome nascent chain complexes (RNC) were generated and prepared as described in previous work^104^. Briefly, protein coding sequences with a N-terminal spytag^105^ were amplified without a stop codon. Ribosomes with a spytag labeled L17 subunit was used during *in vitro* translation system (NEB, E3313S) to generate the stalled RNC. The reaction products were supplemented with spycatcher conjugated double stranded DNA for handle attachment. Optical tweezers force-ramp measurements were carried out with a dual-trap optical tweezers instrument (CTrap, Lumicks) at ambient temperature (22 °C).

## Acknowledgments

This work was supported by a Grant from the National Institutes of Health (5R01GM121567) to CK. We acknowledge Dr. Hao Zhang at the Cell Sorting Core Facility in JHMI for cell sorting operations, Dr. Will Ludington for generously sharing the flow cytometer, Dr. David Mohr at GRCF in JHMI for sequencing operations, and Advanced Research Computing at Hopkins (ARCH) core facility (rockfish.jhu.edu) for high-performance computation resources.

## Supplementary information

**Supp. Figure 1.**
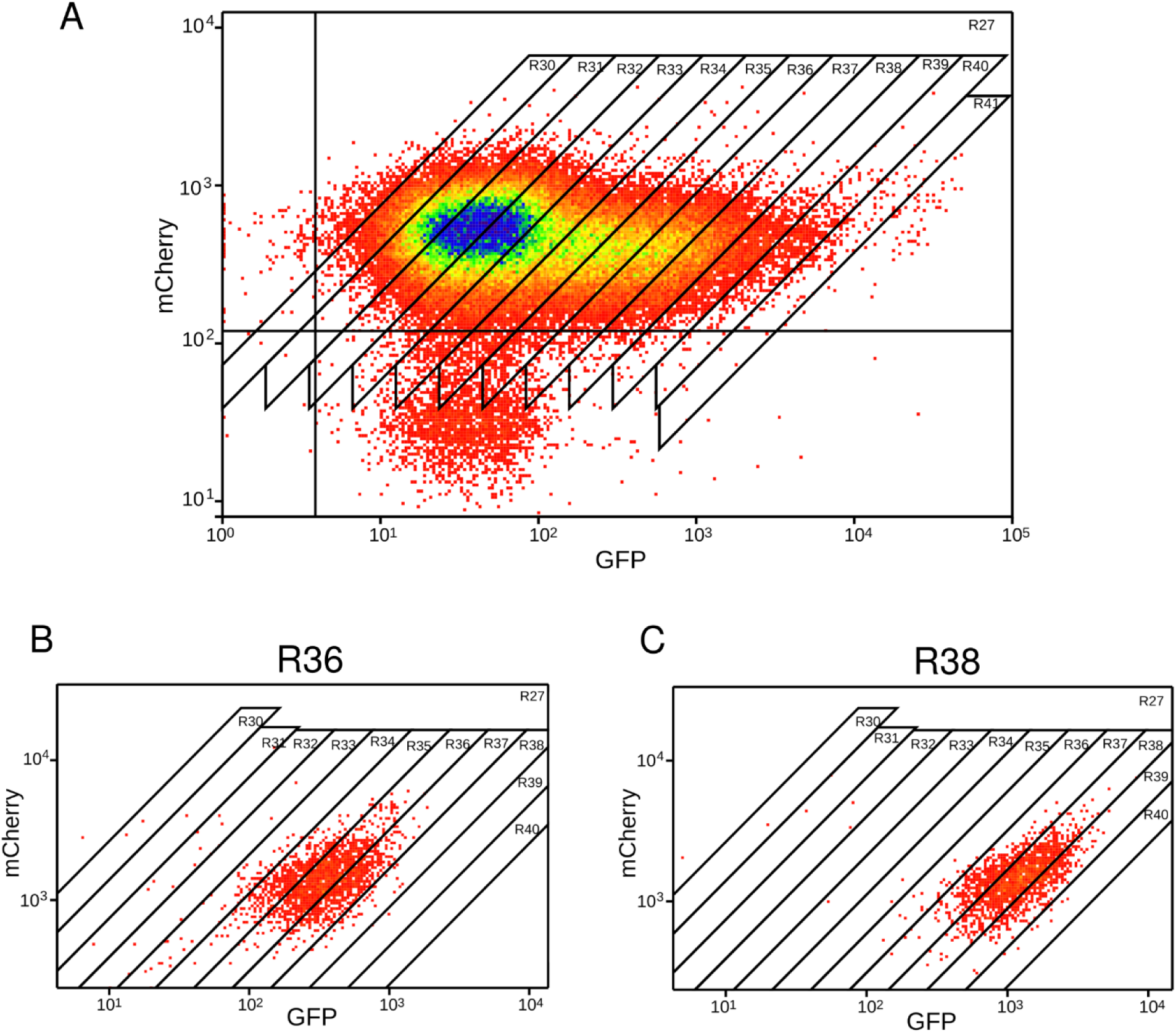
Bacteria sorting purity check. (**A**) An example of quantification sort during AP profiling. R30 - R41 labels the gate settings we used for sorting. The sort logic only included events within the top right quadrant (R27). Sorted samples were injected back to the cytometer for detection and gate settings remained the same. (**B**) The sorted events from R36 reappeared near R36. (**C**) The sorted events from R38 reappeared near R38. The variance in fluorescence detection of particles in the cytometer resulted in slight deviations from the gate boundaries. Plots were cropped directly from the Summit software and enlarged for clarity.

**Supp. Figure 2.**
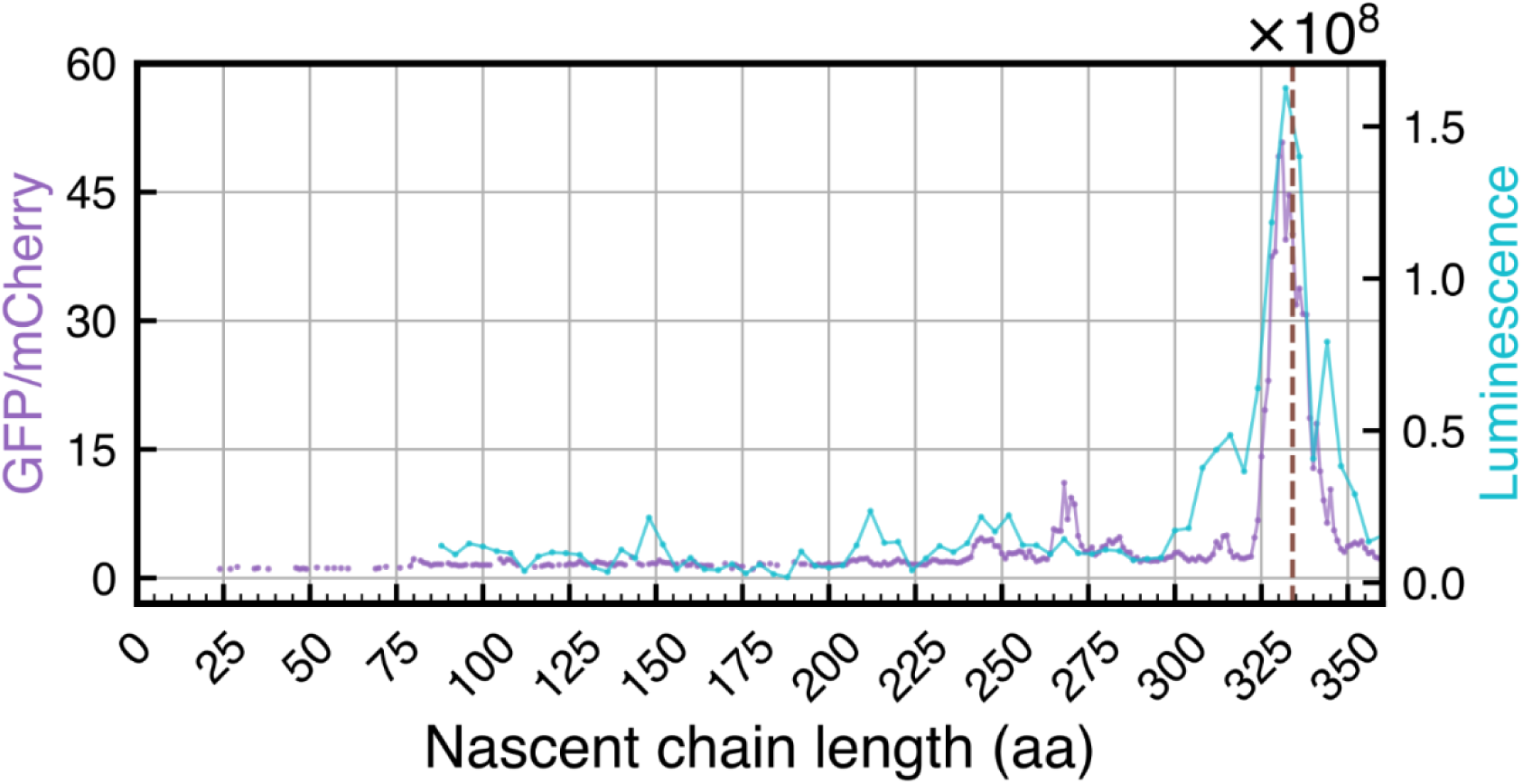
Linear AP profiling results are consistent with previous findings. The peak position and general trend of measurements are consistent from AP profiling and previous work^38^. The dashed line indicates G-domain extrusion.

**Supp. Figure 3.**
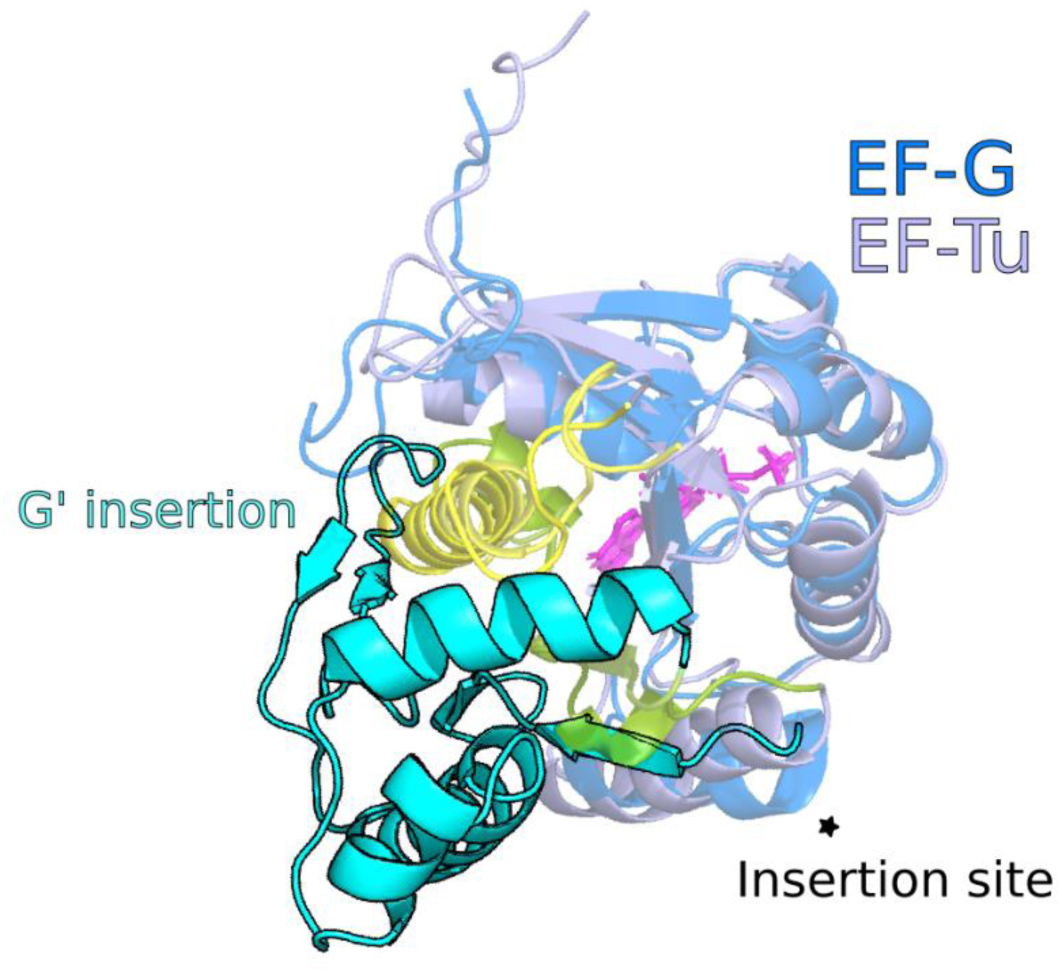
Structure comparison between G_EF-G_ and G_EF-Tu_. The two structures are aligned by the core P-loop domain (residues 1-162 for both proteins, EF-G in blue and EF-Tu in light blue), the C-terminal helices from both proteins are colored yellow. The last strand connecting to the C-terminal helix is colored green. G’ insertion from EF-G is highlighted in cyan. The insertion site is noted with a star.

**Supp. Figure 4.**
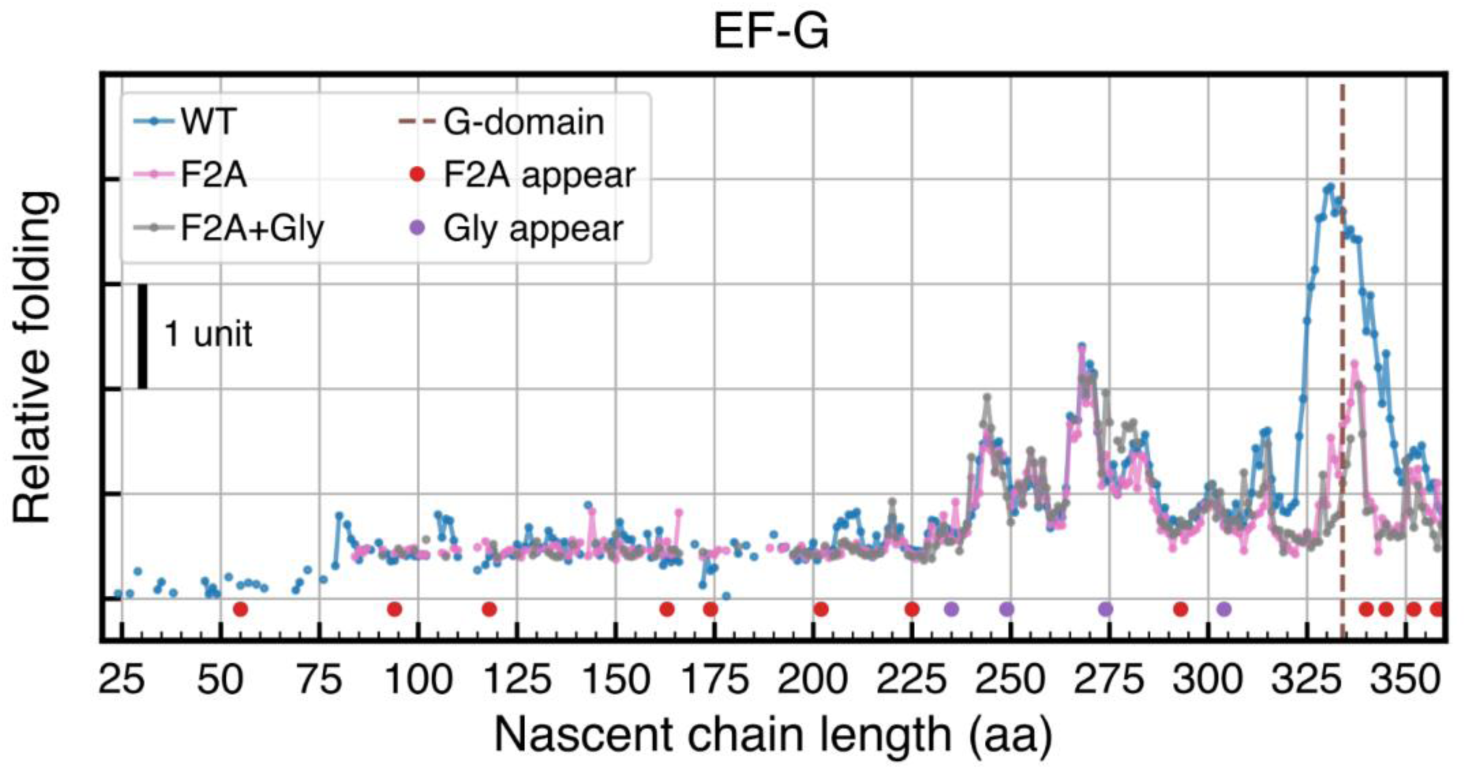
Additional glycine mutations did not significantly disrupt folding peaks. In addition to the F/A mutations (F/A), the indicated residues (V212, A226, A251, A281) were mutated to glycine and subject to AP profiling (F/A+Gly, n=1). WT (n=3) and F2A data (n=2) are the average from repeat experiments.

**Supp. Figure 5.**
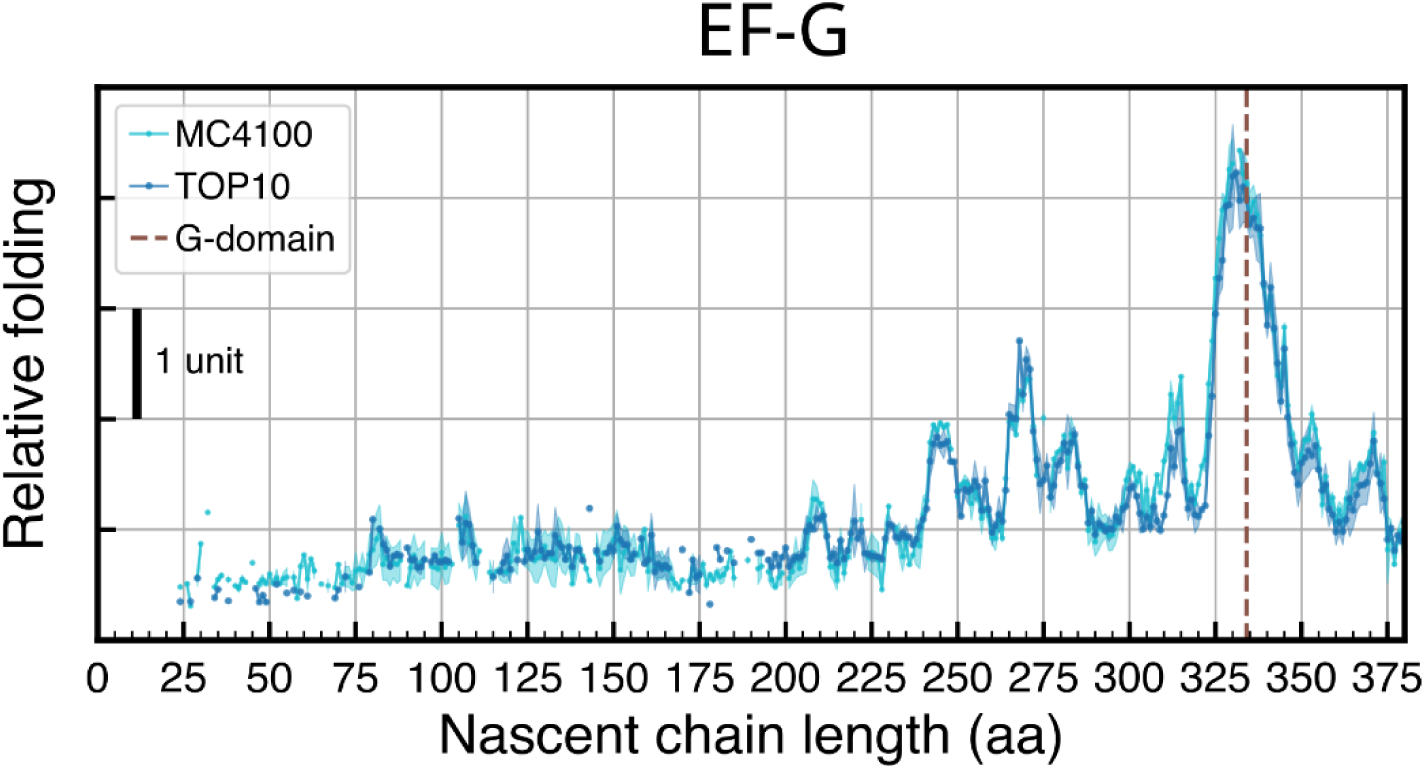
AP profiling results across different *E. coli* strains. Comparison of AP profiling results in different E. coli strains. For each strain, the dots represent the average from 3 repeats and the shaded area is one standard deviation from the average.

